# Characterization of a putative NsrR homologue in *Streptomyces venezuelae* reveals a new member of the Rrf2 superfamily

**DOI:** 10.1101/050989

**Authors:** John T. Munnoch, M^a^ Teresa Pellicer Martinez, Dimitri A. Svistunenko, Jason C. Crack, Nick E. Le Brun, Matthew I. Hutchings

## Abstract

Members of the Rrf2 superfamily of transcription factors are widespread in bacteria but their functions are largely unexplored. The few that have been characterized in detail sense nitric oxide (NsrR), iron limitation (RirA), cysteine availability (CymR) and the iron sulfur (Fe-S) cluster status of the cell (IscR). In this study we combined ChIP- and dRNA-seq with *in vitro* biochemistry to characterize a putative NsrR homologue in *Streptomyces venezuelae.* ChIP-seq analysis revealed that rather than regulating the nitrosative stress response like *Streptomyces coelicolor* NsrR, Sven6563 binds to a conserved motif at a different, much larger set of genes with a diverse range of functions, including a number of regulators, genes required for glutamine synthesis, NADH/NAD(P)H metabolism, as well as general DNA/RNA and amino acid/protein turn over. Our biochemical experiments further show that Sven6563 has a [2Fe-2S] cluster and that the switch between oxidized and reduced cluster controls its DNA binding activity *in vitro.* To our knowledge, both the sensing domain and the putative target genes are novel for an Rrf2 protein, suggesting Sven6563 represents a new member of the Rrf2 superfamily. Given the redox sensitivity of its Fe-S cluster we have tentatively named the protein RsrR for <u>Redox s</u>ensitive response <u>Regulator.</u>

## Introduction

Filamentous *Streptomyces* bacteria produce bioactive secondary metabolites that account for more than half of all known antibiotics as well as anticancer, anti-helminthic and immunosuppressant drugs ^1,2^. More than 600 *Streptomyces* species are known and each encodes between 10 and 50 secondary metabolites but only 25% of these compounds are produced *in vitro.* As a result, there is huge potential for the discovery of new natural products from *Streptomyces* and their close relatives. This is revitalizing research into these bacteria and *Streptomyces venezuelae* has recently emerged as a new model for studying their complex life cycle, in part because of its unusual ability to sporulate to near completion when grown in submerged liquid culture. This means the different tissue types involved in the progression to sporulation can be easily separated and used for tissue specific analyses such as RNA sequencing and chromatin immunoprecipitation and sequencing (RNA- and ChIP-seq) ^3,4^. *Streptomyces* species are complex bacteria that grow like fungi, forming a branching, feeding substrate mycelium in the soil that differentiates upon nutrient stress into reproductive aerial hyphae that undergo cell division to form spores ^5^. Differentiation is closely linked to the production of antibiotics, which are presumed to offer a competitive advantage when nutrients become scarce in the soil.

*Streptomyces* bacteria are well adapted for life in the complex soil environment with more than a quarter of their ~9 Mbp genomes encoding one and two-component signaling pathways that allow them to rapidly sense and respond to changes in their environment ^6^. They are facultative aerobes and have multiple systems for dealing with redox, oxidative and nitrosative stress. Most species can survive for long periods in the absence of O_2_, most likely by respiring nitrate, but the molecular details are not known ^7^. They deal effectively with nitric oxide (NO) generated either endogenously through nitrate respiration ^7^ or in some cases from dedicated bacterial NO synthase (bNOS) enzymes ^8^ or by other NO generating organisms in the soil ^9^. We recently characterized NsrR, which is the major bacterial NO stress sensor, in *Streptomyces coelicolor*(ScNsrR). NsrR is a dimeric Rrf2 family protein with one [4Fe-4S] cluster per monomer that reacts rapidly with up to eight molecules of NO ^10,11^. Nitrosylation of the Fe-S cluster results in derepression of the *nsrR*, *hmpA1* and *hmpA2* genes ^11^, which results in transient expression of HmpA NO dioxygenase enzymes that convert NO to nitrate ^12–14^. The Rrf2 superfamily of bacterial transcription factors is still relatively poorly characterized, but many have C-terminal cysteine residues that are known or predicted to coordinate Fe-S clusters. Other characterized Rrf2 proteins include RirA which senses iron limitation most likely through an Fe-S cluster ^15^ and IscR which senses the Fe-S cluster status of the cell ^16^.

In this work we report the characterization of the *S. venezuelae* Rrf2 protein Sven6563 that is annotated as an NsrR homologue. In fact, it shares only 27% primary sequence identity with ScNsrR and is not genetically linked to an *hmpA* gene (Supplementary Figure S1). We purified the protein from *E. coli* under anaerobic conditions and found that is a dimer with each monomer containing a reduced [2Fe-2S] cluster that is rapidly oxidized but not destroyed by oxygen. Thus, the [2Fe-2S] cofactor is different to the [4Fe-4S] cofactors in the *S. coelicolor* and *Bacillus subtilis* NsrR proteins. The [2Fe-2S] cluster of Sven6563 switches easily between oxidized and reduced states and we provide evidence that this switch controls its DNA binding activity, with holo-RsrR showing highest affinity for DNA in its oxidised state. We have tentatively named the protein RsrR for Redox sensitive response Regulator. ChIP-seq and ChIP-exo analysis allowed us to define the RsrR binding sites on the *S. venezuelae* genome with RsrR binding to class 1 target genes with an 11-3-11bp inverted repeat motif and class 2 target genes with a single repeat or half site. Class 1 target genes suggest a primary role in regulating NADH/NAD(P)H and glutamate/glutamine metabolism rather than nitrosative stress. The *sven6562* gene, which is divergent from *rsrR*, is the most highly induced transcript, up 5.41-fold (log2), in the Δ *rsrR* mutant and encodes a putative NAD(P)+ binding repressor in the NmrA family. Other class 1 target genes are not significantly affected by loss of RsrR suggesting additional levels of regulation, possibly including the divergently expressed Sven6562 (NmrA). Taken together our data suggest that RsrR is a new member of the Rrf2 family and extends the known functions of this superfamily, potentially sensing redox via a [2Fe-2S] cofactor in a mechanism that has only previously been observed in SoxR proteins.

## Results

### Identifying RsrR target genes in *S. venezualae*

We previously reported a highly specialized function for the NO-sensing NsrR protein in *S. coelicolor.* ChIP-seq against a 3xFlag-ScNsrR protein showed that it only regulates three genes, two of which encode NO dioxygenase HmpA enzymes, and the *nsrR* gene itself ^11^. To investigate the function of RsrR, the putative NsrR homologue in *S. venezuelae*, we constructed an *S. venezualae ΔrsrR* mutant expressing an N-terminally 3xFlag-tagged protein and performed ChIPseq against this strain (accession number GSE81073). The sequencing reads from the wild-type (control) sample were subtracted from the experimental sample before ChIP peaks were called (Figure 1a). Using an arbitrary cut-off of ≥500 sequencing reads we identified 117 enriched target sequences (Supplementary data S1). We confirmed these peaks by visual inspection of the data using Integrated Genome Browser ^17^ and used MEME ^18^ to identify a conserved motif in all 117 ChIP peaks (Figure 1b). In 14 of the 117 peaks this motif is present as an inverted 11-3-11 bp repeat, which is characteristic of full-length Rrf2 binding sites^16,19^, and we called these class 1 targets (Figure 1c). In the other 103 peaks it is present as a single motif or half site and we call these class 2 targets (Figure 1b).. The divergent genes *sven3827/8* contain a single class 1 site and the 107 bp intergenic region between *sven6562* and *rsrR* contains two class 1 binding sites separated by a single base pair. It seems likely that RsrR autoregulates and also regulates the divergent *sven6562*, which encodes a LysR family regulator with an NmrA-type ligand-binding domain. These domains are predicted to sense redox poise by binding NAD(P)+ but not NAD(P)H ^20^. The positions of the two RsrR binding sites relative to the transcript start sites (TSS) of *sven6562* and *rsrR* suggests that RsrR represses transcription of both genes by blocking the RNA polymerase binding site (Supplementary Figure S2). Following investigation of RNA-seq expression data (Supplementary data S1) comparing the wild-type and Δ*rsrR* strains the only ChIP-seq associated class 1 target with a significantly altered expression profile is *sven6562* which is ~5.41-fold (log2) induced by loss of RsrR. We hypothesis that other class 1 targets for which we have RNA-seq data are not significantly affected because they are subject to additional levels of regulation, including perhaps by Sven6562 itself although this remains to be seen.

**Figure 1.**
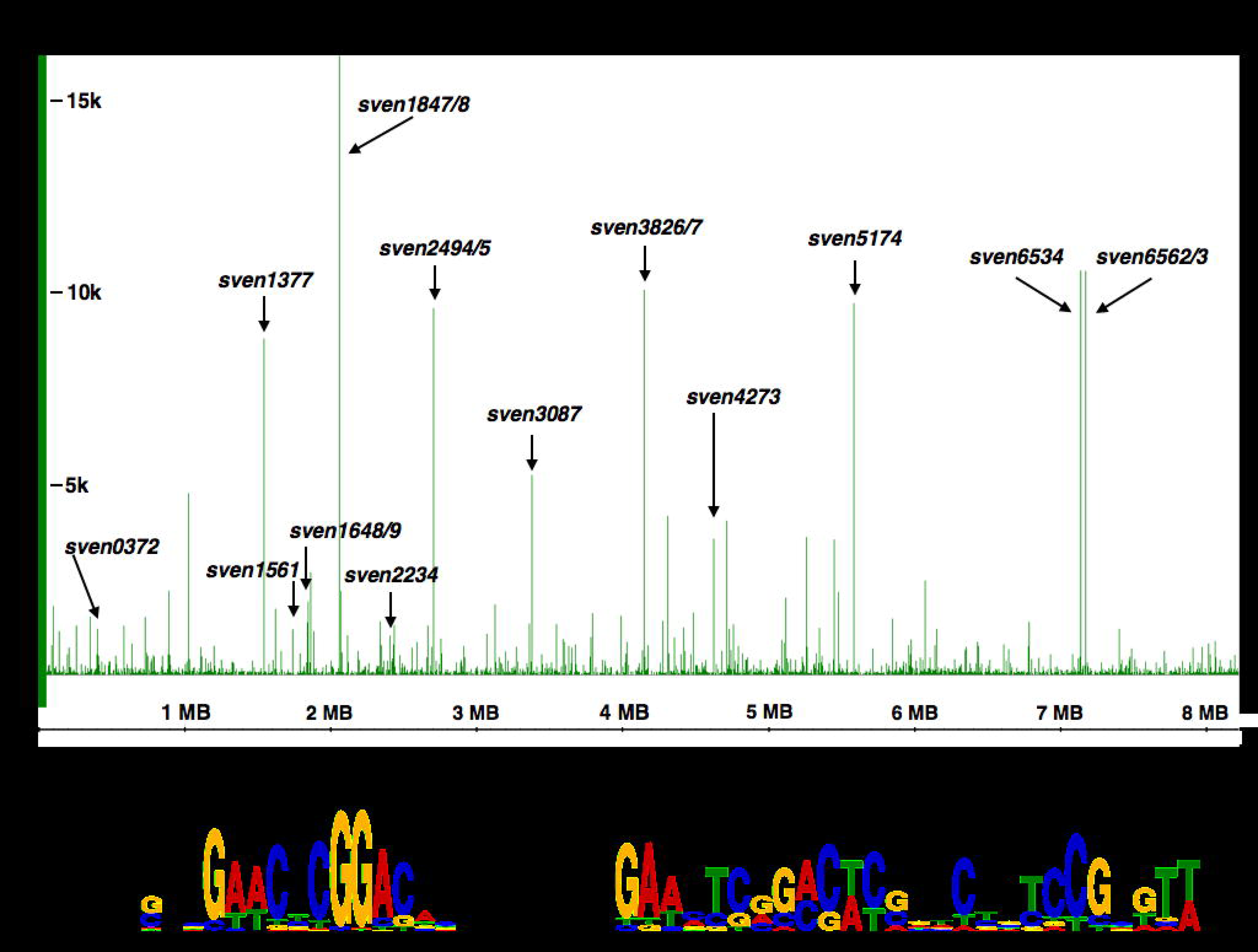
Defining the regulon and binding site for RsrR. Top panel (a) shows the whole genome ChIP-seq analysis with class 1 sites labeled in black. The frequency of each base sequenced is plotted with genomic position on the x-axis and frequency of each base sequenced on the y-axis for *S. venezualae* (NC_018750). Bottom panel (b) shows the class 1 and 2 web logos generated following MEME analysis of the ChIP-seq data.

Other class 1 targets include the *nuo* (NADH dehydrogenase) operon *sven4265-78* (*nuoA-N*) which contains an internal class 1 site upstream of *nuoH,* the putative NADP+ dependent dehydrogenase Sven1847 and the quinone oxidoreductase Sven5173 which converts quinone and NAD(P)H to hydroquinone and NAD(P)+ (Table 1). These data suggest a role for RsrR in regulating NAD(P)H metabolism. In addition to the genes involved directly in NADH/NAD(P)H metabolism, class 2 targets include 21 putative transcriptional regulators, genes involved in both primary and secondary metabolism, RNA/DNA replication and modification genes, transporters (mostly small molecule), proteases, amino acid (particularly glutamate and glutamine) metabolism, and a large number of genes with of unknown function (Supplementary data S1).

**Table 1.** Combined ChIP-Seq and dRNA-Seq data for selected RsrR targets. A – Genes flanking the ChIP-seq peak B – Distance to the translational start codon (bp). C – Distance to the transcriptional start site (bp). D– EMSA shift reactions have been carried out successfully and specifically on these targets E– Relative expression (Log2) fold change WT vs. *RsrR::apr* mutant F– Expression values defined for targets with >100 mapped reads. Class 2 targets are highlighted in red.

| Flanking gene<sup>a</sup> |  |  |  |  |  |  |
| --- | --- | --- | --- | --- | --- | --- |
| Left (-1) | Right (+1) | Distance<sup>b</sup> | Dist. TSS<sup>c</sup> | Fold change<sup>e</sup> | Annotation | Additional description |
|  | *sven0372<sup>d</sup>* | 7 | -99 | -0.73 | Two-component system histidine kinase | Involved in a two-component system signal transduction set |
|  | *sven0519<sup>d</sup>* | -993 |  | 0.53 | Sulfate permease | Involved in sulfate uptake |
|  | *sven0772* | -408 |  | N/A | Xaa-Pro aminopeptidase | Peptidase releasing N-terminal amino acid next to a proline |
|  | *sven1561<sup>d</sup>* | 103 | 36 | -0.11 | Glutamine synthase | Carries out the reaction: Glutamate + NH<sub>4</sub> -> Glutamine |
|  | *sven1670* | 17 |  | -0.28 | Pyridoxamine 5'-phosphate oxidase | Involved in steps of the vitamin B6 metabolism pathway |
| *sven1685* |  | 1215 |  | 0.37 | CoA-transferase family III domain | - |
|  | *sven1847<sup>d</sup>* | 6 |  | -0.89 | 3-oxoacyl-[acyl-carrier protein] reductase | Carries out: NADP+ dependant reduction of 3-oxoacyl-[ACP] |
|  | *sven1902* | -1643 | -1689 | -0.03 | Glutamine synthase adenyltransferase | Regulates glutamine synthase activity by adenylation |
|  | *sven2494* | 90 | 0 | -1.69 | Hypothetical protein | - |
|  | *sven2540* | 221 |  | N/A | Glucose fructose oxidoreductase | D-glucose + D-fructose <> D-gluconolactone + D-glucitol |
|  | *sven3087* | 2092 | 2092 | -0.02 | Acetyltransferase | Transfers an acetyl group |
|  | *sven3827<sup>d</sup>* | 26 | -10 | 0.15 | SAICAR synthetase | Involved in purine metabolism |
| *sven3934* |  | 875 |  | -0.21 | Enhanced intracellular survival protein | - |
|  | *sven4022* | -772 |  | -0.55 | Hypothetical protein | NAD(P)-binding Rossmann-like domain |
| *sven4272* | *sven4273* | 1362, -573 |  | 0.01 | NADH-ubiquinone oxidoreductase chain I | Involved in the electron transfer chain, binds a [4Fe-4S] |
|  | *sven5088* | -77 |  | -0.15 | Epimerase/dehydratase | NADH dependant isomerase enzyme |
|  | *sven5174<sup>d</sup>* | -119 |  | -0.18 | Quinone oxidoreductase | H<sub>2</sub> + menaquinone <> menaquinol |
|  | *sven6227* | -1279 |  | -5.21 | NADH-FMN oxidoreductase | FMNH<sub>2</sub> + NAD<sup>+</sup> <> FMN + NADH + H<sup>+</sup> |
| *sven6534* |  | -100 |  | 0.97 | Trypsin-like peptidase domain | A serine protease that hydrolyses proteins |
| *sven6562<sup>d</sup>* | *sven6563<sup>d</sup>* | 72, -35 | 36 | 5.41<sup>F</sup>, N/A | *nmrA/rsrR* | DNA binding proteins, NADP/[2Fe-2S] binding |

### Purified RsrR contains a redox active [2Fe-2S] cluster

The genes bound by RsrR do not include any NO detoxification genes and this suggested it is not an NsrR homologue but instead has an alternative function. To learn more about the protein we purified it from *E. coli* under strictly anaerobic conditions. The anaerobic RsrR solution is pink in colour but rapidly turns brown when exposed to O_2_, suggesting the presence of a redox-active cofactor. Consistent with this, the UV-visible absorbance spectrum of the as-isolated protein revealed broad weak bands in the 300-640 nm region but following exposure to O_2_, the spectrum changed significantly, with a more intense absorbance band at 460 nm and a pronounced shoulder feature at 330 nm (Figure 2a). The form of the reduced and oxidized spectra are similar to those previously reported for [2Fe-2S] clusters that are coordinated by three Cys residues and one His ^21,22^. The anaerobic addition of dithionite to the previously air-exposed sample (at a 1:1 ratio with [2Fe-2S] cluster as determined by iron content) resulted in a spectrum very similar to that of the as-isolated protein (Figure 2a), demonstrating that the cofactor undergoes redox cycling.

**Figure 2.**
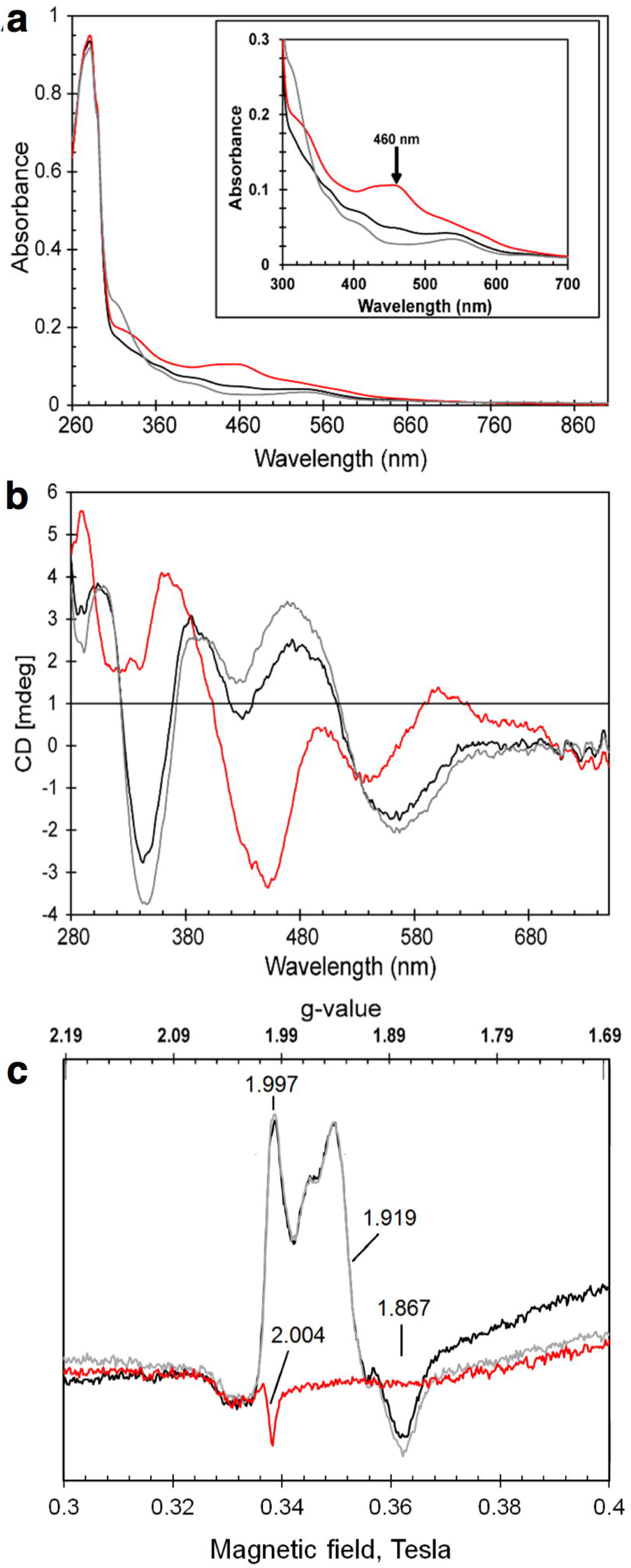
Spectroscopic characterization of RsrR. UV-visible absorption **(a**), CD (**b**) and EPR spectra (c) of 309 µM [2Fe-2S] RsrR (~75% cluster-loaded). Black lines – as isolated, red lines – oxidised, grey lines reduced proteins. In (a) and (b), initial exposure to ambient O_2_ for 30 min was followed by 309 µM sodium dithionite treatment; in (c) – as isolated protein was first anaerobically reduced by 309 µM sodium dithionite and then exposed to ambient O 2 for 50 min. A 1 mm pathlength cuvette was used for optical measurements. Inset in (a) shows details of the iron-sulfur cluster absorbance in the 300 – 700 nm region.

Because the electronic transitions of iron-sulfur clusters become optically active as a result of the fold of the protein in which they are bound, CD spectra reflect the cluster environment ^23^. The near UV-visible CD spectrum of RsrR (Figure 2b) for the as-isolated protein contained three positive (+) features at 303, 385 and 473 nm and negative (-) features at 343 and 559 nm. When the protein was exposed to ambient O_2_ for 30 min, significant changes in the CD spectrum were observed, with features at (+) 290, 365, 500, 600 nm and (-) 320, 450 and 534 nm (Figure 2b). The CD spectra are similar to those reported for Rieske-type [2Fe-2S] clusters ^21,24,25^, which are coordinated by two Cys and two His residues. Anaerobic addition of dithionite (1 equivalent of [2Fe-2S] cluster) resulted in reduction back to the original form (Figure 2b) consistent with the stability of the cofactor to redox cycling.

The absorbance data above indicates that the cofactor is in the reduced state in the as-isolated RsrR protein. [2Fe-2S] clusters in their reduced state are paramagnetic (S = ½) and therefore should give rise to an EPR signal. The EPR spectrum for the as-isolated protein contained signals at g = 1.997, 1.919 and 1.867 (Figure 2c). These g-values and the shape of the spectrum are characteristic of a [2Fe-2S]^1+^ cluster. The addition of excess sodium dithionite to the as-isolated protein did not cause any changes in the EPR spectrum (Figure 2c) indicating that the cluster was fully reduced as isolated. Exposure of the as-isolated protein to ambient O_2_ resulted in an EPR-silent form, with only a small free radical signal typical for background spectra, consistent with the oxidation of the cluster to the [2Fe-2S]^2+^ form (Figure 2c), and the same result was obtained upon addition of the oxidant potassium ferricyanide (data not shown).

To further establish the cofactor that RsrR binds, native ESI-MS was employed. Here, a C-terminal His-tagged form of the protein was ionized in a volatile aqueous buffered solution that enabled it to remain folded with its cofactor bound. The deconvoluted mass spectrum contained several peaks in regions that corresponded to monomer and dimeric forms of the protein, (Supplementary Figure S3). In the monomer region (Figure 3a), a peak was observed at 17,363 Da, which corresponds to the apo-protein (predicted mass 17363.99 Da), along with adduct peaks at +23 and +64 Da due to Na^+^(commonly observed in native mass spectra) and most likely two additional sulfurs (Cys residues readily pick up additional sulfurs as persulfides, respectively ^26^. A peak was also observed at +176 Da, corresponding to the protein containing a [2Fe-2S] cluster. As for the apo-protein, peaks corresponding to Na^+^and sulfur adducts of the cluster species were also observed (Figure 3a). A significant peak was also detected at +120 Da that corresponds to a break down product of the [2Fe-2S] cluster (from which one iron is missing, FeS_2_). In the dimer region, the signal to noise is significantly reduced but peaks are still clearly present (Figure 3b). The peak at 34,726 Da corresponds to the RsrR homodimer (predicted mass 34727.98 Da), and the peak at +352 Da corresponds to the dimer with two [2Fe-2S] clusters.

**Figure 3.**
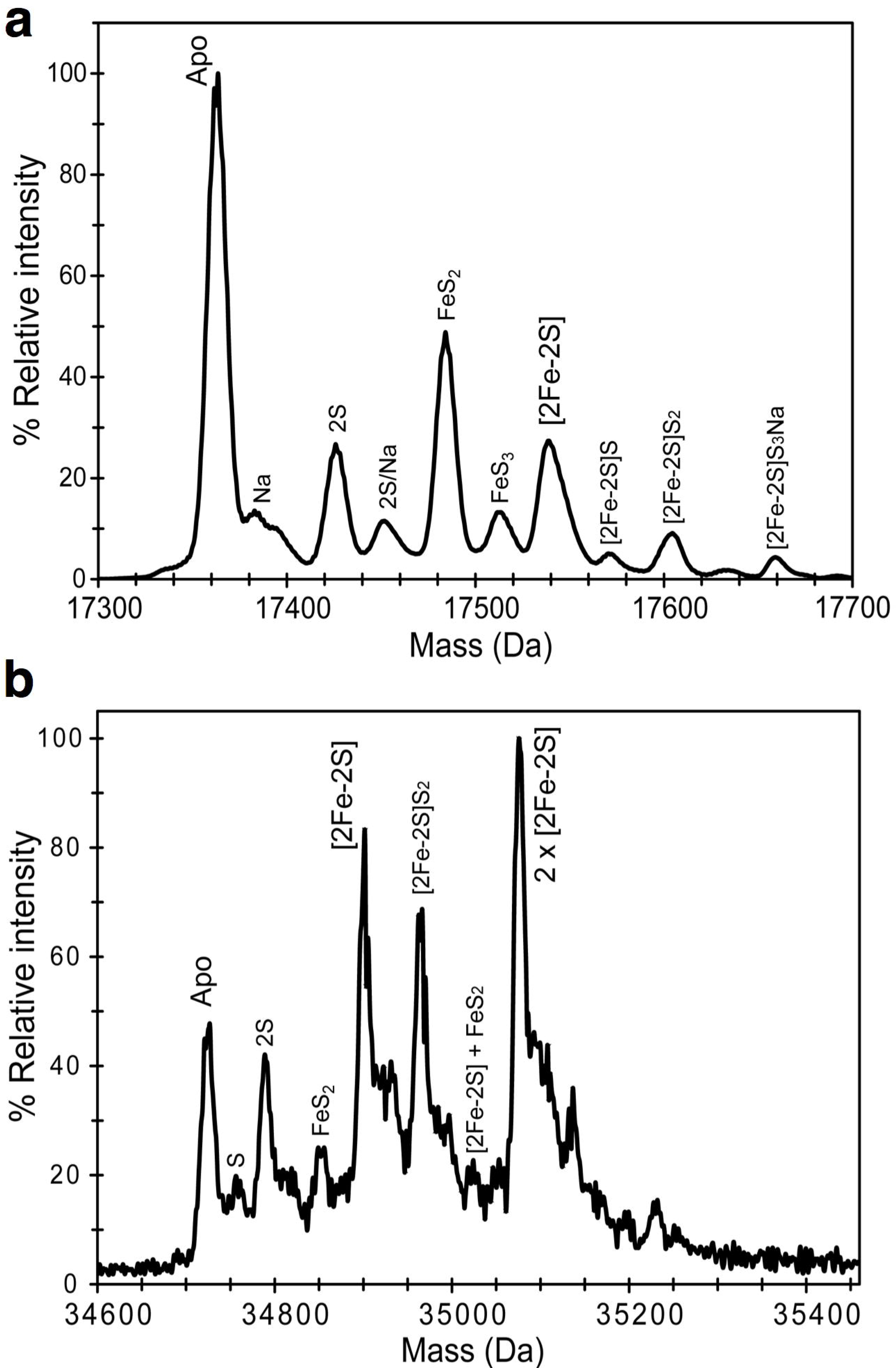
Native mass spectrometry of RsrR. (**a**) and (**b**) Positive ion mode ESI-TOF native mass spectrum of ~21 µM [2Fe-2S] RsrR in 250 mM ammonium acetate pH 8.0, in the RsrR monomer (a) and dimer (b) regions. Full *m*/*z* spectra were deconvoluted with Bruker Compass Data analysis with the Maximum Entropy plugin.

A peak at +176 Da is due to the dimer containing one [2Fe-2S] cluster. A range of cluster breakdown products similar to those detected in the monomer region were also observed (Figure 3b). Taken together, the data reported here demonstrate that RsrR contains a [2Fe-2S] cluster that can be reversibly cycled between oxidised (+2) and reduced (+1) states.

### Cluster and oxidation state dependent binding of RsrR *in vitro*

To determine which forms of RsrR are able to bind DNA, we performed EMSA experiments using the intergenic region between the highly enriched ChIP target *sven1847/8* as a probe. Increasing ratios of [2Fe-2S] RsrR to DNA resulted in a clear shift in the mobility of the DNA from unbound to bound, see Figure 4a. Equivalent experiments with cluster-free (apo) RsrR did not result in a mobility shift, demonstrating that the cluster is required for DNA-binding activity. These experiments were performed aerobically and so the [2Fe-2S] cofactor was in its oxidised state. To determine if oxidation state affects DNA binding activity, EMSA experiments were performed with [2Fe-2S]^2+^ and [2Fe-2S]^1+^ forms of RsrR. The oxidised cluster was generated by exposure to air and confirmed by UV-visible absorbance. The reduced cluster was obtained by reduction with sodium dithionite, confirmed by UV-visible absorbance, and the reduced state was maintained using EMSA running buffer containing an excess of dithionite. The resulting EMSAs, Figure 4b and c, show that DNA-binding occurred in both cases but that the oxidised form bound significantly more tightly. Tight binding could be restored to the reduced RsrR samples by allowing it to re-oxidise in air (data not shown). We cannot rule out that the apparent low affinity DNA binding observed for the reduced sample results from partial re-oxidation of the cluster during the electrophoretic experiment. Nevertheless, the conclusion is unaffected: oxidised, [2Fe-2S]^2+^ RsrR is the high affinity DNA-binding form and these results suggest a change in the redox state of the [2Fe-2S] cluster controls the activity of RsrR, something which has only previously been observed for SoxR, a member of the MerR superfamily ^27^.

**Figure 4.**
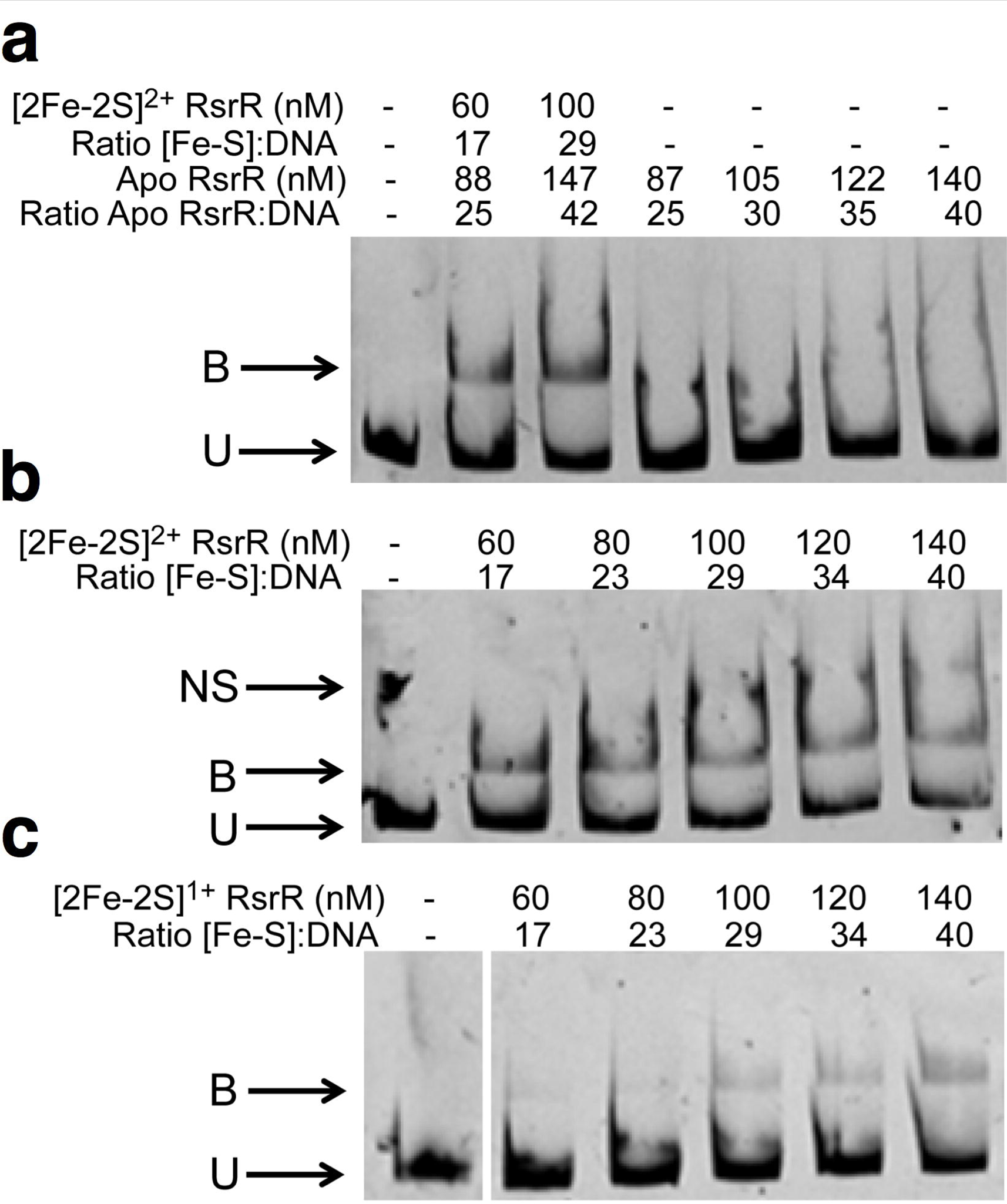
Cluster- and oxidation state-dependent DNA binding by [2Fe-2S] RsrR. EMSAs showing DNA probes unbound (U), bound (B), and non-specifically bound (NS) by (**a**) [2Fe-2S]^2+^ and apo-RsrR (**b**) [2Fe-2S]^2+^ RsrR and (**c**) [2Fe-2S]^1+^ RsrR. Ratios of [2Fe-2S] containing RsrR (Holo) and [RsrR] (apo) to DNA are indicated for (a) while the concentration of [2Fe-2S] RsrR only is reported in (b) and (c). DNA concentration was 3.5 nM for the [2Fe-2S]^2+/1+^ and apo-RsrR experiments. For (a) and (b) the reaction mixtures were separated at 30 mA for 50 min and the polyacrylamide gels were pre-run at 30 mA for 2 min prior to use. For (c) the reaction mixtures were separated at 30 mA for 1h 45 min and the polyacrylamide gel was pre-run at 30 mA for 50 min prior to use using the de-gassed running buffer containing 5 mM sodium dithionite. For (a) both holo and apo protein concentrations are represented as the sample contained both forms due to incomplete cluster loading. The concentrations reported are of the [2Fe-2S] concentration.

### Oxidised [2Fe-2S] RsrR binds strongly to class 1 and 2 binding sites *in vitro*

To further investigate the DNA binding activities of [2Fe-2S]^2+^ RsrR, EMSAs were performed on DNA probes containing the two class 2 RsrR binding sites at *sven0247* and *sven519* (Figure 5a). Both probes were shifted by oxidized [2Fe-2S] RsrR showing that RsrR binds to both class 1 and 2 probes *in vitro*. To further test the idea of RsrR recognizing full and half sites, we constructed a series of probes based on the divergent *nmrA*-*rsrR* promoters carrying both or each individual natural class 1 sites (Figure 5b) and artificial half sites (Figure 5c). The combinations of artificial half sites are illustrated in Supplemental Figure S3 in regards to the original promoter region. The results show that RsrR binds strongly to both full class 1 binding sites at the *nmrA-rsrR* promoters (Figure 5b) but only weakly to artificial half sites (Figure 5c). This suggests that although MEME only calls half sites in most of the RsrR target genes identified by ChIP-seq these class 2 targets must contain sufficient sequence information in the other half to enable strong binding by RsrR.

**Figure 5.**

Oxidised RsrR binding to full site (class 1) and half site (class 2) RsrR targets. EMSAs showing DNA probes unbound (U) and bound (B) by [2Fe-2S]^2+^. Ratios of [2Fe-2S] RsrR and [RsrR] to DNA are indicated. DNA concentration was 4 nM for each probe. EMSA’s using class 2 DNA probes *sven0247* and *sven0519*(a), class 1 probes from the RsrR *rsrR* binding region (b) and the four possible half sites from the *rsrR* class 1 sites (c) were used. For (a) the reaction mixtures were separated at 30 mA for 1h and the polyacrylamide gel was pre-run at 30 mA for 2 min prior to use. For (b) and (c) the reaction mixtures were separated at 30 mA for 30 min and the polyacrylamide gels were prerun at 30 mA for 2 min prior to use. A representation of the *rsrR* promoter breakdown is also available in Supplementary Figure S3b.

### Mapping RsrR binding sites *in vivo* using ChIP-exo and differential RNA-seq

MEME analysis of the ChIP-seq data detected only 14 class 1 (11-3-11bp inverted repeat) sites out of the 117 target sites bound by RsrR on the *S. venezuelae* chromosome. However, ChIP-Seq and EMSAs show that RsrR can bind to target genes whether they contain class 1 or class 2 sites. This differs from *E. coli* NsrR which binds only weakly to target sites containing putative half sites (class 2) ^28^. To gain more information about RsrR recognition sequences and the positions of these binding sites at target promoters we combined differential RNA-seq (dRNA-seq, accession number GSE81104), which maps the start sites of all expressed transcripts, with ChIP-exo (accession number GSE80818) which uses Lambda exonuclease to trim excess DNA away from ChIP complexes leaving only the DNA which is actually bound and protected by RsrR. For dRNA-seq, total RNA was prepared from cultures of wild type *S. venezuelae* and for the Δ*rsrR* mutant grown for 16 hours. ChIP-exo was performed on the Δ*rsrR* strain producing Flag-tagged RsrR, also at 16 hours. ChIP-exo identified 630 binding sites which included the 117 targets identified previously using ChIP-seq. The ChIP-exo peaks are on average only ~50bp wide giving much better resolution of the RsrR binding sites at each target. MEME analysis using all 630 ChIP-exo sequences identified the class 2 binding motif in every sequence and we identified transcript start sites (TSS) for 261 of the 630 RsrR target genes using our dRNA-seq data (Supplementary data S1). Figure 6 shows a graphical representation of class 1 targets that have clearly defined TSS, indicating the centre of the ChIP peak, the associated TSS and any genes within the ~200 bp frame. Based on the RsrR binding site position at putative target genes RsrR likely acts as both a transcriptional activator and repressor and we have shown that RsrR represses transcription of *sven6562* which is a class 1 target with two 11-3-11bp binding site in the intergenic region between *sven6562* and *rsrR.* The functional significance of RsrR binding to the other class 1 and 2 target genes identified here by ChIP-seq and ChIP-exo remains to be seen but they are not significantly affected by loss of RsrR under the conditions used in our experiments.

**Figure 6.**

Graphical representation of combined ChIP-Seq, ChIP-exo and dRNA-seq for four class 1 targets. Each target has the relative position of ChIP-exo (blue line) peak centre (dotted line) and putative transcriptional start site (TSS - solid arrow) indicated with the distance in bp (black numbers) relative to the down stream start codon of target genes. The y-axis scale corresponds to number of reads for ChIP data with each window corresponding to 200 bp with each ChIP-peak being ~50 bp wide. Above each is the relative binding site sequence coloured following the weblogo scheme (A – red, T – green, C – blue and G – yellow) from the MEME results.

## Discussion

In this work we have identified and characterized a new member of the Rrf2 protein family, which was mis-annotated as an NsrR homologue in the *S. venezuelae* genome. ChIP analyses show that RsrR binds to 630 sites on the *S. venezuelae* genome which compares to just three target sites for *S. coelicolor* NsrR and their DNA recognition sequences are very different. RNA-seq data shows a dramatic 60-fold change in the expression of the divergent gene from *rsrR*, *sven6562*, but under normal laboratory conditions no other direct RsrR targets are significantly induced or repressed by loss of RsrR. Approximately 2.7% of the RsrR targets contain class 1 binding sites which consist of a MEME identified 11-3-11 bp inverted repeat. Class 1 target genes include *sven6562* and are involved in either signal transduction and / or NAD(P)H metabolism which perhaps points to a link to redox poise and recycling of NAD(P)H to NAD(P)*in vivo*. The >600 class 2 target genes contain only half sites with a single repeat but exhibit strong binding by RsrR *in vitro.* Our EMSA experiments show that RsrR binds weakly to artificial half sites and this suggests additional sequence information is present at class 2 binding sites that increases the strength of DNA binding by RsrR. Six of the class 2 targets are involved in glutamate and glutamine metabolism including: *sven1561*, encoding a Glutamine Synthase (GS) that carries out the ATP dependent conversion of glutamate and ammonium to glutamine^29^, *sven1902*, encoding a GS adenylyltransferase that carries out the adenylation and deadenylation of GS, reducing or increasing GS activity respectively^30^. *sven3711,* encoding a protein which results in the liberation of glutamate from glutamine^31^. *sven4418,* encoding a glutamine fructose-6-phosphate transaminase that carries out the reaction: L-glutamine and D-fructose 6-phosphate to L-glutamate and D-glucosamine 6-phosphate^32^. *sven4888,* encoding a glutamate-1-semialdehyde aminotransferase, which carries out the PLP dependent, reversible reaction of L-glutamate to 1-semialdehyde 5-aminolevulinate^33^. Finally, *sven7195*, encoding a glutamine-dependent asparagine synthase which carries out the ATP dependent transfer of NH_3_ from glutamine to aspartate, forming glutamate and asparagine^34^. Glutamate and glutamine are precursors for the production of mycothiol, the actinobacterial equivalent of glutathione, which acts as a cellular reducing agent. Mycothiol also acts as a cellular reserve of cysteine and in the detoxification of redox species and antibiotics^35^. Glutamate is important, as a non-essential amino acid, because it links nitrogen and carbon metabolism in bacteria^36^. Additionally, glutamate acts as a proton sink through its decarboxylation to GABA, which especially under acidic conditions, favorably removes protons from the intracellular milieu^37^.

Our data show that the purified RsrR protein contains a [2Fe-2S] cluster, which is stable in the presence of O_2_ and can be reversibly cycled between reduced (+1) and oxidized (+2) states. The oxidised [2Fe-2S]^2+^ form binds strongly to both class 1 and class 2 binding sequences *in vitro*, whereas the reduced [2Fe-2S]^1+^ form exhibits significantly weaker binding. The binding we did observed is likely due to partial oxidation of the RsrR Fe-S cluster during the EMSA electrophoresis. The cluster free form of RsrR does not bind to DNA at all. Given these observations and the stability of the Fe-S cluster to aerobic conditions, we propose that the activity of RsrR is modulated by the oxidation state of its cluster, becoming activated for DNA binding through oxidation and inactivated through reduction. Exposure to O_2_ is sufficient to cause oxidation, but other oxidants may also be important *in vivo*. The properties of RsrR described here are reminiscent of the *E. coli*[2Fe-2S] cluster containing transcription factor SoxR, which controls the regulation of another regulator, SoxS, through the oxidation state of its cluster^38^.

Due to the number of RsrR regulated transcription factors it is likely that its target genes are subject to multiple levels of regulation. For example, the *sven6562* gene, which is divergent from *rsrR*, encodes a LysR family regulator with an N terminal NmrA-type NAD(P)+ binding domain. NmrA proteins are thought to control redox poise in fungi by sensing the levels of NAD(P), which they can bind, and NAD(P)H, which they cannot^39^. This is intriguing and we propose a model in which reduction of holo-RsrR induces expression of Sven6562 which in turn senses redox poise via the ratio of NAD(P)/NAD(P)H and then modulates expression of its own regulon which likely overlaps with that of RsrR. Clearly there is much to learn about this system and it will be important to define the role of Sven6562 in *S. venezuelae* in the future. We did not observe any phenotype for the Δ*rsrR* mutant and it is no more sensitive to redox active compounds or oxidative stress than wild-type *S. venezuelae* (not shown). However, this is not surprising given the number of systems in bacteria that deal with reactive nitrogen and oxygen species and redox stress. In *Streptomyces* species these include catalases, peroxidases^40^ and superoxide dismutases^41^ and associated regulators such as OxyR^42^, SigR^43^, OhrR^44^, Rex^20^ and SoxR^45^. Thus, our data suggests Sven6563, tentatively renamed here as RsrR, is a new member of the Rrf2 family and this work extends our knowledge about this neglected but widespread superfamily of bacterial transcription factors.

## Materials and methods

### Bacterial strains, plasmids, oligonucleotides and growth conditions

Bacterial strains and plasmids are listed in Table 2 and oligonucleotides are listed in Table 3. For ChIP-seq experiments, *S. venezuelae* strains were grown at 30°C in MYM liquid sporulation medium^46^ made with 50% tap water and supplemented with 200μl trace element solution^47^ per 100ml and adjusted to a final pH.of 7.3. Disruption of *rsrR* was carried out following the PCR-targeting method^48^ as described previously^49,50^. Primers JM0109 and JM0110 were used to PCR amplify the apramycin disruption cassette from pIJ773. Cosmid SV-5-F05 was used as the template cosmid. The disruption cosmid (pJM026) was checked by PCR using primers JM0111 and JM0112. Antibiotic marked, double crossover exconjugants, were identified as previously described and confirmed once more with JM0111 and JM0112. The 3x Flag tag copy of *rsrR* was synthesized by Genescript and subcloned into pMS82 using HindIII/KpnI and confirmed by PCR using primers JM0113 and JM0114.

**Table 2.** Strains and plasmids used during this study.

| Strain/plasmid | Description | Source |
| --- | --- | --- |
| *E. coli* |  |  |
| TOP10 | F<sup>-</sup> mcrA Δ(mrr-hsdRMS-mcrBC) φ80lacZΔM15 ΔlacX74 nupG recA1 araD139 Δ(ara-leu)7697 galE15 galK16 rpsL(Str<sup>R</sup>) endA1 λ<sup>-</sup> | Invitrogen |
| BW25113 (pIJ790) | *E. coli* BW25113 containing λRED recombination plasmid pIJ790 | 48,58 |
| ET12567 (pUZ8002) | *E. coli* Δdam dcm strain containing helper plasmid pUZ8002 | 59,60 |
| BL21 | F<sup>-</sup> ompT gal dcm lon hsdS<sub>B</sub>(r<sub>B</sub><sup>-</sup> m<sub>B</sub><sup>-</sup>) λ(DE3 [lacI lacUV5-T7 gene 1 ind1 sam7 nin5]) | 61 |
| *Streptomyces* |  |  |
| *S. venezuelae* | *S. venezuelae* ATCC 10712 WT strain | 4 |
| *rsrR::apr* | *S. venezuelae* with a ReDirect disrupted *sven6563::apr* |  |
| *rsrR::apr 3xFlag RsrR* | *rsrR::apr* with a pMS82 encoded N-terminal 3xFlag tagged *rsrR* with 300 bp of upstream flanking DNA (promoter) | This work |
| Plasmids |  |  |
| pIJ773 | pBluescript KS (+), aac(3)IV, oriT (RK2), FRT sites | 48 |
| SV-5-F05 | Supercos-1-cosmid with (a 52181 bp) fragment containing *sven6562/3* | 4 |
| pMS82 | *ori*, pUC18, *hyg*, *oriT*, RK2, int ΦBT1 | 62 |
| pGS-21a | Genscript overexpression and purification vector (SD0121) | Genscript |
| pJM026 | SV-5-F05 containing *sven6563::apr oriT* | This work |
| pJM027 | pMS82, *rsrR* gene plus 300 bp upstream DNA with a c-terminal synthetic linker and 3xFLAG tag | This work |
| pJM028 | pGS-21a, full length *rsrR* cloned NdeI/XhoI | This work |
| pJM029 | pJM028 with a c-terminal 6xHis tag NdeI/XhoI | This work |
| pJM030 | pJM028 with a c-terminal synthetic linker as with (flag), 2xFLAG tag and a 6xHis tag, cloned NdeI/XhoI | This work |

**Table 3.**
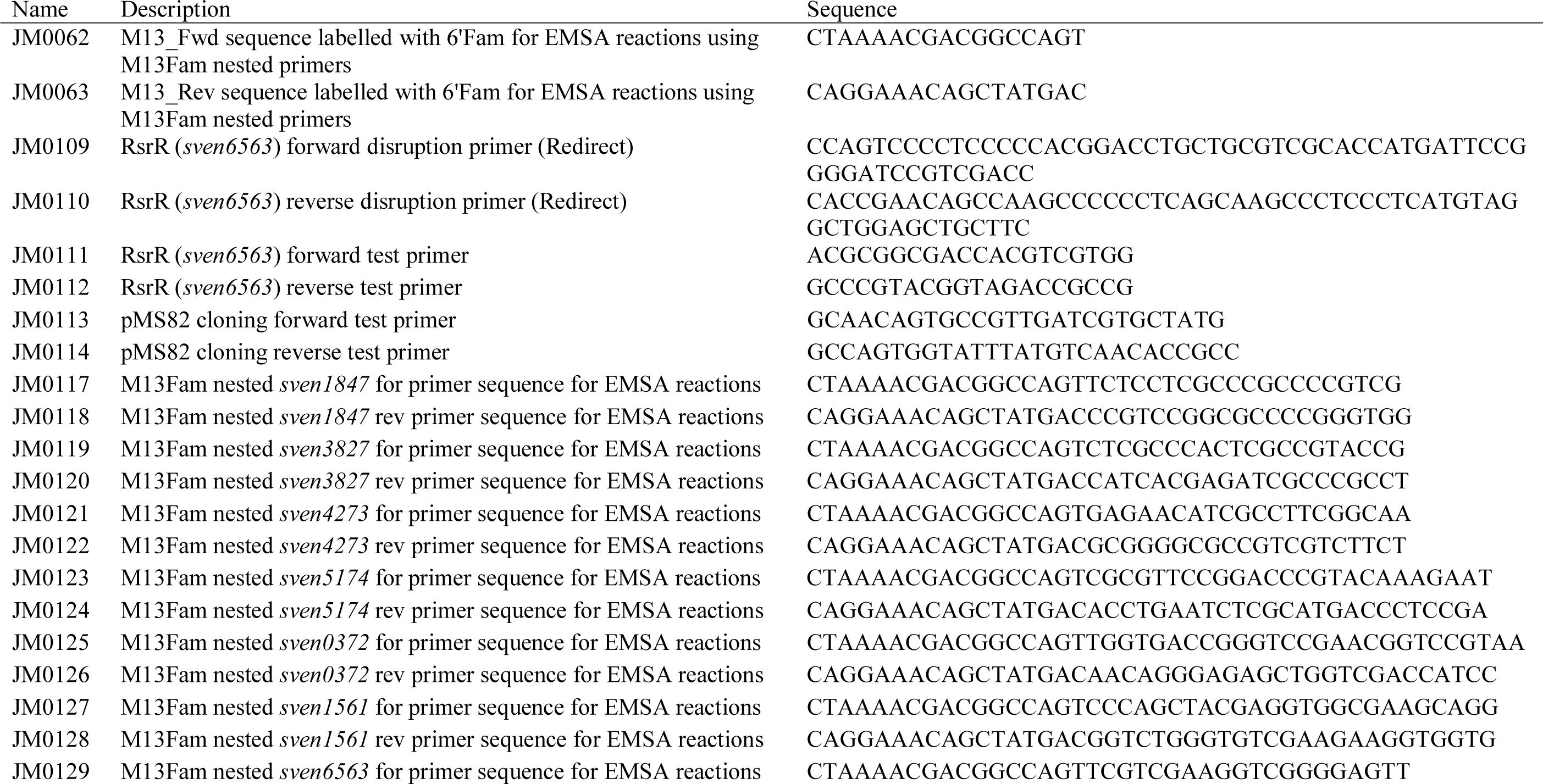
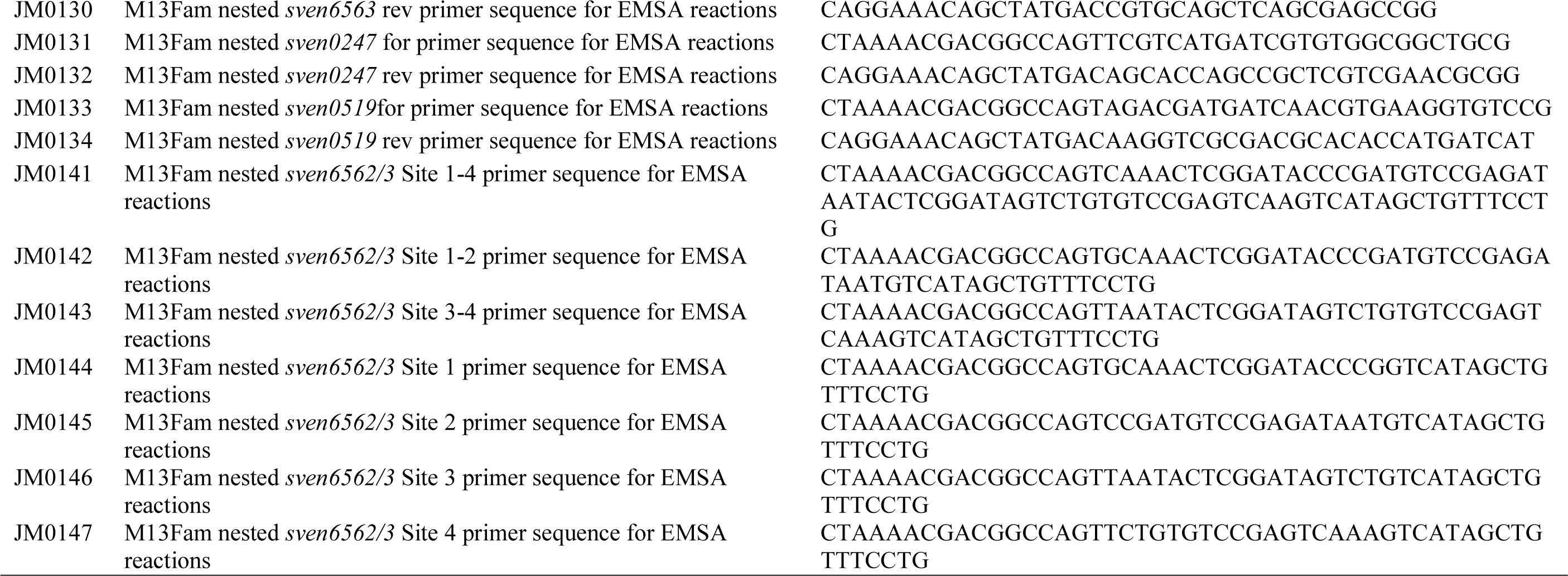
List of primers used in this study. Primers JM0119-JM0134 were used to produce EMSA DNA templates that were successfully shifted using purified RsrR and mentioned in the text but the data is not shown as part of the work.

### ChIP (chromatin immunoprecipitation) – seq and exo

ChIP-Seq was carried out as previously described^51^ with the below modifications. A 3xFlag tagged RsrR was used as with our previous work^11^. Following sonication and lysate clearing M2 affinity beads (Sigma-Aldrich #A2220) were prepared by washing in ½IP buffer following manufacturers instructions. The cleared lysate was incubated with 40 µl of washed M2 beads and incubated for 4 h at 4C in a vertical rotor. The lysate was removed and the beads pooled into one 1.5 microfuge tube and washed in 0.5 IP buffer. The beads were transferred to a fresh microfuge tube and washed a further 3 times removing as much buffer as possible without disturbing the beads. The DNA-protein complex was eluted from the beads with 100 µl elution buffer (50 mM Tris-HCl pH7.6, 10mM EDTA, 1% SDS) by incubating at 65°C overnight. Removing the ~100µl elution buffer, an extra 50 µl of elution buffer was added and further incubated at 65°C for 5 min. To extract the DNA 150 µl eluate, 2 µl proteinase K (10 mg/ml) was added and incubated 1.5 h at 55°C. To the reaction 150 µl phenol-chloroform was added. Samples were vortexed and centrifuged at full speed for 10 min. The aqueous layer was extracted and purified using the Qiaquick column from Qiagen with a final elution using 50 µl EB buffer (Qiagen). The concentration of samples were determined using Quant-iT™ PicoGreen^®^ dsDNA Reagent (Invitrogen) or equivalent kit or by nanodrop measurement.

DNA sequencing of ChIP-Seq samples was carried out by GATC. ChIP-exo following sonication of lysates was carried out by Peconic LLC (State College, PA) adding an additional exonuclease treatment to the process as previously described^52^.

Data analysis was carried out using CLC workbench 8 followed by a manual visual inspection of the data.

### dRNA - seq

Mycelium was harvested at experimentally appropriate time points and immediately transferred to 2 ml round bottom tubes, flash frozen in liquid N2, stored at −80°C or used immediately. All apparatus used was treated with RNaseZAP (Sigma) to remove RNases for a minimum of 1 h before use. RNaseZAP treated mortar and pestles were used, the pestle being placed and cooled on a mixture of dry ice and liquid N_2_ with liquid N_2_ being poured into the bowl and over the mortar. Once the bowl had cooled the mycelium samples were added directly to the liquid N_2_ and thoroughly crushed using the mortar leaving a fine powder of mycelium. Grindings were transferred to a pre-cooled 50 ml Falcon tube and stored on dry ice. Directly to the tube, 2 ml of TRI reagent (Sigma) was added to the grindings and mixed. Samples are then thawed while vortexing intermittently at room temperature for 5-10 min until the solution cleared. To 1 ml of TRI reagent resuspension, 200 μl of chloroform was added and vortexed for 15 seconds at room temperature then centrifuged for 10 min at 13,000 rpm. The upper, aqueous phase (clear colourless layer) was removed into a new 2 ml tube. The remainder of the isolation protocol follows the RNeazy Mini Kit (Qiagen) instructions carrying out both on and off column DNase treatments. On column treatments were carried out following the first RW1 column wash. DNaseI (Qiagen) was added (10 *μ*l enzyme, 70 *μ*l RDD buffer) to the column and stored at RT for 1 h. The column was washed again with RW1 then treated as described in the manufacturer’s instructions. Once eluted from the column, samples were treated using TURBO DNA-free Kit (Ambion) following manufacturer’s instructions to remove residual DNA contamination.

Data analysis was carried out using the Tuxedo protocal^53^ for analysis of gene expression and TSSAR webservice for dRNA transcription start site analysis^54^. In addition a manual visual processing approach was carried out for each.

### Purification of RsrR

5 L Luria-Bertani medium (10 × 500 mL) was inoculated with freshly transformed BL21 (DE3) *E. coli* containing a pGS-21a vector with the *prsrR-His* insert. 100 µg/mL ampicillin and 20 µM ammonium ferric citrate were added and the cultures were grown at 37 °C, 200 rpm until OD_600 nm_ was 0.6-0.9. To facilitate *in vivo* iron-sulfur cluster formation, the flasks were placed on ice for 18 min, then induced with 100 µM IPTG and incubated at 30 °C and 105 rpm. After 50 min, the cultures were supplemented with 200 µM ammonium ferric citrate and 25 µM L-Methionine and incubated for a further 3.5 h at 30 °C. The cells were harvested by centrifugation at 10000 × g for 15 min at 4 °C. Unless otherwise stated, all subsequent purification steps were performed under anaerobic conditions inside an anaerobic cabinet (O_2_ < 2 ppm). Cells pellets were resuspended in 70 mL of buffer A (50 mM TRIS, 50 mM CaCl_2_, 5% (v/v) glycerol, pH 8) and placed in a 100 mL beaker. 30 mg/mL of lysozyme and 30 mg/mL of PMSF were added and the cell suspension thoroughly homogenized by syringe, removed from the anaerobic cabinet, sonicated twice while on ice, and returned to the anaerobic cabinet. The cell suspension was transferred to O-ring sealed centrifuge tubes (Nalgene) and centrifuged outside of the cabinet at 40,000 × g for 45 min at 1 °C.

The supernatant was passed through a HiTrap IMAC HP (1 × 5mL; GE Healthcare) column using an ÄKTA Prime system at 1 mL/min. The column was washed with Buffer A until A_280 nm_ <0.1. Bound proteins were eluted using a 100 mL linear gradient from 0 to 100% Buffer B (50 mM TRIS, 100 mM CaCl_2_, 200mM L-Cysteine, 5% glycerol, pH 8). A HiTrap Heparin (1 × 1mL; GE Healthcare) column was used to remove the L-Cysteine, using buffer C (50 mM TRIS, 2 M NaCl, 5% glycerol, pH 8) to elute the protein. Fractions containing RsrR-His were pooled and stored in an anaerobic freezer until needed. RsrR-His protein concentrations were determined using the method of Bradford (BioRad Laboratories)^55^, with BSA as the standard. Cluster concentrations were determined by iron assay^56^, from which an extinction coefficient, ε, at 455 nm was determined as 3450 ±25 M-1 cm-1, consistent with values reported for [2Fe-2S] clusters with His coordination^21^.

### Preparation of Apo-RsrR

Apo-RsrR -His was prepared from as isolated holoprotein by aerobic incubation with 1 mM EDTA overnight.

### Spectroscopy and mass spectrometry

UV-visible absorbance measurements were performed using a Jasco V500 spectrometer, and CD spectra were measured with a Jasco J810 spectropolarimeter. EPR measurements were performed at 10 K using a Bruker EMX EPR spectrometer (X-band) equipped with a liquid helium system (Oxford Instruments). Spin concentrations in the protein samples were estimated by double integration of EPR spectra with reference to a 1 mM Cu(II) in 10 mM EDTA standard. For native MS analysis, His-tagged RsrR was exchanged into 250 mM ammonium acetate, pH 8, using PD10 desalting columns (GE Life Sciences), diluted to ~21 µM cluster and infused directly (0.3 mL/h) into the ESI source of a Bruker micrOTOF-QIII mass spectrometer (Bruker Daltonics, Coventry, UK) operating in the positive ion mode. Full mass spectra (*m*/*z* 700–3500) were recorded for 5 min. Spectra were combined, processed using the ESI Compass version 1.3 Maximum Entropy deconvolution routine in Bruker Compass Data analysis version 4.1 (Bruker Daltonik, Bremen, Germany). The mass spectrometer was calibrated with ESI-L low concentration tuning mix in the positive ion mode (Agilent Technologies, San Diego, CA).

### Electrophoretic Mobility Shift Assays (EMSAs)

DNA fragments carrying the intergenic region between *sven1847* and *sven1848 of the S. venezualae* chromosome were PCR amplified using *S. venezualae* genomic DNA with 5’ 6-FAM modified primers (Table 2). The PCR products were extracted and purified using a QIAquick gel extraction kit (Qiagen) according to the manufacturer’s instructions. Probes were quantitated using a NanoDrop ND2000c. The molecular weights of the double stranded FAM labelled probes were calculated using OligoCalc^57^.

Bandshift reactions (20 µl) were carried out on ice in 10 mM Tris, 60 mM KCl, pH 7.52. Briefly, 1 µL of DNA was titrated with varying aliquots of RsrR. 2 µL of loading dye (containing 0.01% (w/v) bromophenol blue), was added and the reaction mixtures were immediately separated at 30 mA on a 5% (w/v) polyacrylamide gel in 1 X TBE (89 mM Tris,89 mM boric acid, 2 mM EDTA), using a Mini Protean III system (Bio-Rad). Gels were visualized (excitation, 488 nm; emission, 530 nm) on a molecular imager FX Pro (Bio-Rad). Polyacrylamide gels were pre-run at 30 mA for 2 min prior to use. For investigations of [2Fe-2S]^1+^ RsrR DNA binding, in order to maintain the cluster in the reduced state, 5 mM of sodium dithionite was added to the isolated protein and the running buffer (de-gassed for 50 min prior to running the gel). Analysis by UV-visible spectroscopy confirmed that the cluster remained reduced under these conditions.

## Acknowledgements

We are grateful to the Natural Environment Research Council for a PhD studentship to John Munnoch, to the Biotechnology and Biological Sciences Research Council for the award of grant BB/J003247/1 (to NLB and MIH), to the UEA Science Faculty for a PhD studentship to Maria Teresa Pellicer Martinez. The funders had no role in study design, data collection and interpretation, or the decision to submit the work for publication. We are grateful to Dr Govind Chandra at the John Innes Centre for advice about ChIP- and dRNA-seq data analysis and to UEA for supporting the mass spectrometry facility. The research presented in this paper was carried out on the High Performance Computing Cluster supported by the Research and Specialist Computing Support service at the University of East Anglia. All sequence data was deposited online with the Geo superSeries accession number GSE81105 (ChIP-Seq, ChP-exo and dRNA-seq all at a 16 h time point with accession numbers GSE81073, GSE80818, and GSE81104 respectively).

## Conflict of interest

The authors declare that they have no conflicts of interest with the contents of this article.

## Author contributions

JTM carried out all of the molecular microbiology experiments, some of the biochemical experiments, analysed the data and co-wrote the manuscript. MTPC carried out the bulk of the biochemical experiments, analysed the data and co-wrote the manuscript. JCC analyzed data and co-wrote the manuscript. DAS performed EPR experiments and analysed data. NLB and MIH conceived and coordinated the study, analyzed data and co-wrote the manuscript.

